# SST-MAE: Learning Spectral-Spatio-Temporal Representations from Plant Hyperspectral Time Series to Discover Complex Genotype-Phenotype Relations

**DOI:** 10.64898/2026.07.11.737920

**Authors:** Frank G. Okyere, Sarah L. Mehrem, Basten L. Snoek, Guido Van den Ackerveken, Sanne Abeln

## Abstract

Understanding the link between genetic variation and observable traits is key to crop breeding. Hyperspectral imaging captures physiological and biochemical profiles, but current supervised methods require costly trait annotations and treat each observation as a static snapshot, ignoring the temporal dynamics of plant development. We introduce SST-MAE, a self-supervised framework that learns genotype-discriminative representations from plant hyperspectral developmental trajectories, without requiring phenotypic labels. The model learns to reconstruct masked information, capturing multiple growth trajectories. Validated on 194 field-grown lettuce genotypes across eight time points, the frozen encoder serves as a feature extractor for downstream genotype classification. SST-MAE outperforms raw spectral and linear baselines, achieving AUROC > 0.89 for anthocyanin pigmentation SNPs and 0.77 for leaf serration. The learned features are highly label-efficient, attaining near-full performance with only 30–50% of labeled data, offering a scalable pathway toward high-throughput genetic screening from image-based phenotypes.

## 1 Introduction

Understanding the relationship between genetic variation and observable traits is fundamental to modern crop breeding. Functional single-nucleotide polymorphisms (SNPs) are key markers linked to agronomic traits such as yield and stress tolerance [1]. Genome-wide association studies (GWAS) can identify candidate SNPs through statistical association with phenotypes. However, traditional phenotyping is labor-intensive, destructive, and difficult to scale. Moreover, phenotypic expression is dynamic and often manifests through subtle physiological changes throughout plant development [2], yet most current approaches treat plant observations as static snapshots, failing to capture the temporal dynamics through which many genetic effects gradually emerge.

Hyperspectral imaging (HSI) provides rich physiological, biochemical, and structural information [3]. SNP variants alter plant biochemistry, leaving measurable signatures in HSI data, raising the question: can plant genotypes be inferred directly from hyperspectral phenotypes? Success would enable highthroughput genetic screening from image-based data. While deep learning has advanced HSI analysis for trait prediction and stress detection [4], direct genotype inference remains underexplored. Existing methods require costly, manually annotated trait labels [5, 6] and ignore developmental time-series [7].

Self-supervised learning (SSL), particularly Masked Autoencoders (MAEs), reduces reliance on labels by learning from unlabeled data. Extensions to HSI such as SatMAE [7] and SpectralMAE [17] incorporate spatial and spectral masking but lack temporal modeling. VideoMAE [9] models time but operates on RGB data, ignoring the rich spectral dimension. Plant HSI has unique properties: spectral bands are highly correlated, successive time points share similar information, and subtle genetic shifts become apparent only across full developmental trajectories. Existing methods fail to jointly model the spatial, spectral, and temporal axes, missing the critical interplay through which genetic variation influences not only static morphology but also the timing and rate of phenotypic change. Large labeled HSI datasets with paired genomic information remain extremely rare in plant phenotyping due to the cost of repeated field imaging and sequencing. SSL is thus well suited here as it maximizes unlabeled observations while minimizing dependence on expensive genotype annotations. Our objective is not to compete with foundation models trained on millions of images, but to improve representation learning in realistic breeding datasets.

To address this, we introduce the first self-supervised framework that learns genotype-discriminative representations from plant hyperspectral developmental trajectories. Our approach, SST-MAE, achieves this through joint spatial, spectral, and temporal masking within a Vision Transformer, forcing the model to capture the temporal dynamics through which genetic effects manifest. The model learns latent representations by reconstructing masked information, capturing developmental trajectories rather than static snapshots. After pre-training, the frozen encoder serves as a label-efficient feature extractor for downstream binary SNP classification.

Our key contributions are: (1) the first self-supervised framework that learns genotype-discriminative representations from plant hyperspectral developmental trajectories; (2) a scalable, transferable encoder that enables multi-SNP prediction with limited labeled data; and (3) temporal representation learning enabling developmental trajectory modeling, few-shot prediction, cross-time transfer, and time-to-detection of genetic effects. We validate SST-MAE on a field-grown lettuce panel of 194 genotypes observed across eight time points spanning multiple developmental stages.

## 2 Related Works

Traditional genotype-phenotype association methods rely on linear or Bayesian models [10]. More recent deep learning approaches have employed CNNs, transformers, and attention mechanisms [11]. Stylianou et al. [12] showed CNNs can predict SNP genotypes from sorghum imagery, and subsequent works extended this to HSI for variety-level classification across quinoa, maize, and sorghum [13]. Deep autoencoders have mapped visual crop architecture to haplotype blocks and QTLs [14]. Despite these advances, all such approaches are static, failing to exploit temporal dynamics.

Self-supervised learning (SSL) has transformed visual representation learning. MAE [8] and BEiT [16] established masked reconstruction as a powerful pretraining paradigm. Extensions to HSI, such as SatMAE [7] and Spectral-MAE [17], incorporate spatial and spectral masking but are designed for static observations. While VideoMAE [9] and TimeSformer [18] demonstrate temporal masking for RGB video, they lack spectral depth.

To our knowledge, SST-MAE is the first framework for joint spatial, spectral, and temporal masked representation learning in plant HSI time-series. This gap is critical because genetic variation influences not only static spectra and morphology, but also the timing and rate of phenotypic change throughout development.

## 3 Methodology

### 3.1 Dataset and Preprocessing

#### Plant material and imaging

A lettuce (*Lactuca sativa*) panel of 194 accessions was grown under field conditions in a complete randomised block design with two replicate plots per genotype [2, 5]. Hyperspectral images were acquired at eight time points (57–106 days after sowing) using a Specim FX10 camera (400–1000 nm, 224 bands; ∼1 mm/pixel). Images were calibrated to reflectance using white reference and dark current subtraction, followed by Savitzky–Golay spectral smoothing. Plants were segmented via NDVI thresholding (> 0.3) with morphological refinement and cropped to 128 × 128 pixel bounding boxes.

#### SNP genotyping and selection

Genotype data were obtained from Wei et al. [19], supplemented with 67 additional sequenced lines [20]. Four SNPs with validated candidate genes and significant GWAS associations were selected: ant.loc5 (Chr5, 85.5 Mbp; LsMYB113; anthocyanin), ant.loc9 (Chr9, 152.7 Mbp; LsANS; anthocyanin), pale.loc4 (Chr4, 101.3 Mbp; LsGLK; pale coloration), and serr.loc5 (Chr5, ∼252.7 Mbp; LsTCP4; leaf serration) [2, 5]. We selected four biologically validated SNPs spanning pigmentation and morphology — distinct mechanisms with varying prediction difficulty — to evaluate representation quality rather than maximize prediction targets.

### 3.2 Spatio-Spectro-Temporal Masked Autoencoder (SST-MAE)

To predict SNP genotypes from HSI data, we developed a self-supervised model that takes a sequence of tokenized plant frames as input and learns to reconstruct randomly masked spatial tokens, spectral bands, and entire time frames using a vision transformer (ViT) based encoder-decoder architecture (**Fig. 1**).

**Fig. 1:**
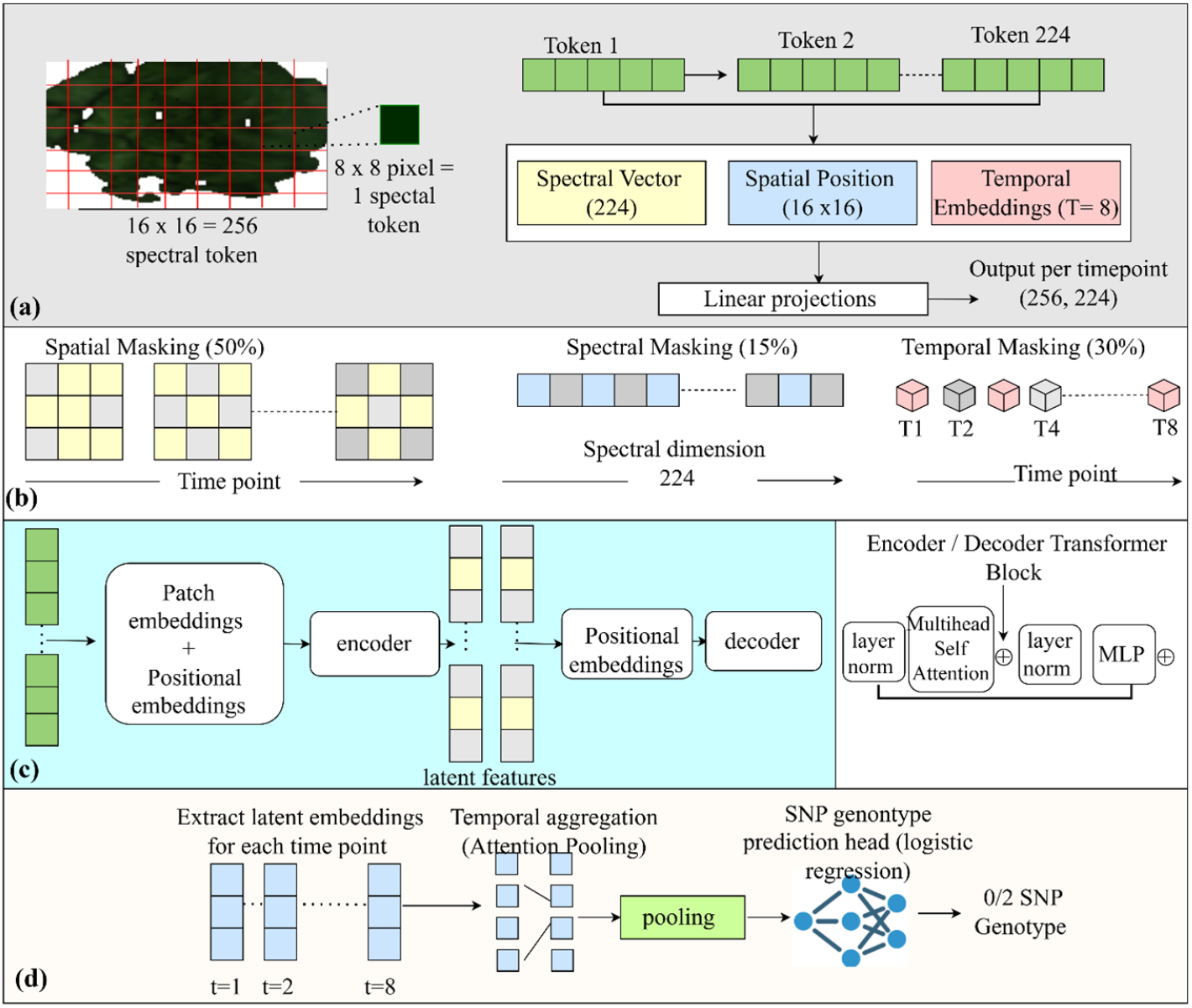
Overview of the SST-MAE workflow. (a) Tokenization: each plant image is partitioned into 8 × 8 px patches (16 × 16 = 256 tokens), each carrying a 224-band spectrum with spatial and temporal positional embeddings. (b) Joint masking: spatial (50%), spectral (15%), and temporal (30%). (c) Encoder-decoder with Pre-layer normalization and residual connections. (d) Latent embeddings are mean-pooled across time and fed to a logistic regression head for binary SNP prediction (0 = reference; 2 = alternate).

#### Input Representation and Positional Embeddings

Each segmented plant image (128 × 128 px) is partitioned into a 16 × 16 grid of 8 × 8 px patches, yielding *N* = 256 spatial tokens per frame. Each token is the mean 224-band spectrum across its pixels. A plant observed at *T* time points is represented as **X** ∈ ℝ^*T* ×*N* ×*D*^ (*D* = 224), linearly projected to embedding dimension *C* = 512: **Z** = **XW**_proj_, where **W**_proj_ ∈ ℝ^*D*×*C*^ .

Two learnable positional embeddings are added to encode spatial and temporal structure:

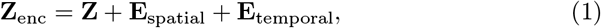

where **E**_spatial_ ∈ ℝ^1×*N* ×*C*^ encodes the 256 spatial positions (shared across time) and **E**_temporal_ ∈ ℝ^*T* ×1×*C*^ encodes developmental progression, with both added via broadcasting. The resulting tensor is flattened to ℝ ^(*TN*)×*C*^ for the transformer encoder.

#### Joint Spatio-Spectro-Temporal Masking

Existing masked autoencoders treat masking dimensions independently; combining SpectralMAE and Video-MAE ignores the biological coupling inherent in plant development—gradual temporal changes and correlated spectral bands. SST-MAE addresses this through simultaneous spatial, spectral, and temporal masking with temporal consistency regularization, explicitly encoding developmental trajectories rather than isolated observations.

We implement three masking strategies applied jointly per batch element:

- *Spectral masking (15%):* Randomly sets 15% of the 224 spectral bands to zero across the entire tensor. This is applied directly to the raw input data to encourage the model to reconstruct fine-grained missing spectral profiles.
- *Temporal masking (30%):* Randomly selects 30% of the *T* developmental time frames to be completely masked. For these omitted time frames, the raw spectral data for all *N* patches within the frame are replaced with a learnable temporal mask tensor prior to the linear projection layer.
- *Spatial masking (50%):* Following linear projection and positional embedding addition, 50% of the spatial tokens within each time frame are randomly masked. The features at these masked locations are replaced with a learnable spatial mask token before the sequence is flattened and fed into the transformer encoder.

All masks are generated independently per batch element.

#### Encoder and decoder Encoder

We use a Vision Transformer Small (ViT Small) [21] with 3 transformer layers (4 heads, key dimension 32, MLP hidden 512, ReLU, dropout 0.1). We chose this lightweight design over deeper (6- and 12-layer) variants because preliminary experiments showed no downstream improvement while increasing training time and overfitting risk—a consequence of our limited dataset (194 genotypes across 8 time points). The selected architecture offers the best trade-off between representation quality, generalization, and resource efficiency.

##### Decoder

The decoder is a lightweight transformer with 2 layers, 4 heads, and an embedding dimension of 128. It takes the encoder output and reconstructs the full sequence of tokens. A final linear layer *W*_out_ ∈ ℝ^*C*×224^ followed by a sigmoid activation maps each token back to 224 spectral bands, producing the reconstruction *X*^*′*^ ∈ ℝ^*B*×*T* ×*N* ×224^. The sigmoid ensures outputs are in the (0, 1) range, matching the normalized input.

#### Loss Function

The total pre-training loss combines spectral reconstruction and temporal regularisation:

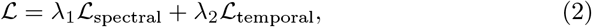

with *λ*_1_ = 1.0 and *λ*_2_ = 0.5 determined by grid search.

The spectral loss is MSE computed exclusively on masked spectral bands:

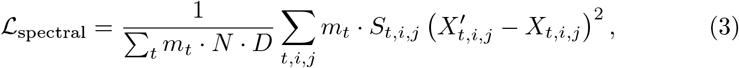

where *i* indexes spatial tokens (*N* = 256), *j* spectral bands (*D* = 224), *t* time frames, *m*_*t*_ excludes padded frames, and *S*_*t,i,j*_ restricts to masked bands.

The temporal loss penalizes large changes in latent space between consecutive frames, enforcing smooth developmental transitions:

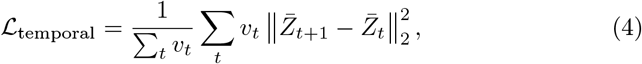

where 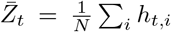 is the spatially averaged latent representation, and *v*_*t*_ confirms both frames are real and unmasked.

#### Training Details

Pre-training used a non-overlapping 80/20 genotype-level split, ensuring both replicate plots of each genotype were assigned exclusively to either partition. The 80% split (11,376 plants) served as the pre-training set; the remaining 20% (2,880 plants) was used solely for monitoring reconstruction loss, learning-rate scheduling, and early stopping. Genotypes in the pre-training partition were withheld from all downstream evaluation (Section 3.3), ensuring the frozen encoder was never optimised on test genotypes. Class distributions per SNP and label fraction are provided in Supplementary Tables S1 and S2.

Input tensors were normalised to [0, 1] using a global maximum computed from training data. The model was optimised using AdamW (lr = 1 × 10^−3^, *β*_1_ = 0.9, *β*_2_ = 0.999, weight decay = 0.05), with learning rate reduced by 0.5 after five epochs without validation improvement and early stopping (patience = 15; maximum 100 epochs). Pre-training performance was assessed using masked-token MSE. The framework was implemented in TensorFlow 2.10+. Computational cost details are provided in Supplementary Table S3.

### 3.3 Downstream Genotype Prediction

#### Frozen feature strategy

For each plant sequence *X*, the data was processed through the encoder to extract frame-wise latent vectors. Spatial representation was handled by computing the mean over all *N* spatial patch tokens within each individual time step, producing a distinct frame-level latent vector *z*_*t*_ ∈ ℝ^*C*^ (where *C* = 128). To generate a single plant-level representation *z*, these vectors were aggregated across the entire observation window by mean pooling over the temporal dimension:

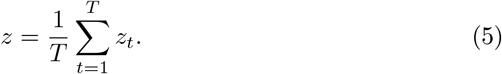

These latent features were then partitioned using the same genotype-level partition described in Section 3.2 (Training Details): the downstream classifier was trained on latent features from the 80% pre-training pool and evaluated on latent features from the 20% of genotypes held out entirely from pre-training. Reusing this partition, rather than drawing an independent random split at the downstream stage, ensures the test genotypes were never seen by the encoder in any capacity—including weight updates, early stopping, or learning-rate scheduling. Within this constraint, the split was additionally stratified where possible to balance the SNP class distribution between the train and test partitions. For each individual SNP, a binary Logistic Regression (LR) classifier was trained using the extracted plant-level feature vectors, with *L*_2_ regularization and balanced class weights. LR model was selected because it is the standard linear evaluation protocol used in self-supervised learning [8, 22]. Linear probing evaluates the intrinsic quality of frozen representations while minimizing the influence of downstream model capacity. Strong performance under a linear classifier shows that the encoder has learned discriminative latent features rather than relying on a powerful task-specific classifier. Only samples with homozygous genotypes (0 = reference, 2 = alternate) were retained; heterozygotes were excluded due to their low frequency and ambiguous spectral presentation. Classification performance was evaluated on the held-out test split using AUROC and macro-F1 score.

### 3.4 Baseline Comparisons

The performance of the SST-MAE model was compared against three baselines:

- **Raw Spectral Mean + Logistic Regression:** A baseline that collapses the data by averaging all raw spectral signatures over all tokens and time points to determine if deep representation extraction is necessary.
- **PCA + Logistic Regression:** A linear baseline consisting of a Principal Component Analysis projection applied directly to the flattened raw token spectra, followed by a downstream Logistic Regression classifier.

All benchmark models were validated using identical plant-level train/test partition boundaries to ensure a fair evaluation.

### 3.5 Temporal Representation Analysis

#### Latent Trajectories

Frame-wise feature vectors were extracted for each plant sequence and projected down to a 2D space utilizing PCA and t-SNE algorithms. Trajectories were visualized by mapping data points along a continuous color gradient reflecting developmental time points, as well as distinct categorical flags highlighting SNP genotypes (0 or 2). This visualization allowed the analysis of genotype-dependent growth patterns and variations in hyperspectral features over time.

#### Time-to-Detection Analysis

For each SNP, Logistic Regression classifiers were trained and evaluated incrementally across developmental stages. At each sequence window *k* (*k* = 1, 2, …, *T*), the features were extracted by taking the mean average of the latent representations over the first *k* time steps. Both the training and testing matrices were restricted strictly to this *k*-frame temporal subset to preserve shape alignment.

#### Cross-Time Generalization

To evaluate temporal generalization properties, genotype classifiers were trained on feature subsets extracted exclusively from the first three chronological time steps (representing early growth) and evaluated using features from the final two time points (representing late growth). This configuration tests whether the features extracted during early developmental stages capture stable, invariant genotypic hallmarks that persist throughout the life cycle of the plant.

#### Ablation Study

To isolate the contribution of each masking dimension within the joint framework, we pre-trained four variants of SST-MAE: (i) the Full model with all three masking modes active, (ii) NoSpatial, where spatial masking was omitted while spectral and temporal masking were retained, (iii) NoSpectral, where spectral masking was omitted while spatial and temporal masking were retained, and (iv) NoTemporal, where temporal masking was omitted while spatial and spectral masking were retained. Following pre-training, features were extracted to measure downstream SNP classification performance (reported as mean AUC across functional SNPs). This ablation isolates the contribution of each masking component.

## 4 Results and Discussion

### 4.1 Pre-training Reconstruction and Unsupervised Latent Topology

To verify that the SST-MAE learns meaningful representations, we evaluated spatial, spectral, and temporal reconstruction quality on the pre-training validation set (defined in Section 3.2, Training Details). For the spectral reconstruction (**Fig. 2a**), the model recovers the overall spectral profile with a remarkably low MSE of 0.0007 across all masked bands, confirming that the spectral branch captures rich biochemical information. The spatial reconstructed heatmap (**Fig. 2b**) closely matches the original canvas, demonstrating the model’s ability to fill missing spatial tokens in the 16 × 16 grid. For this specific evaluation band, the spatial reconstruction achieves an MSE of 0.002, a Peak Signal-to-Noise Ratio (PSNR) of 37.1 dB, and a Structural Similarity Index (SSIM) of 0.913. The temporal reconstruction curve (**Fig. 2c**) tracks true developmental dynamics with minor errors, yielding an MSE of 0.0002. Together, these results demonstrate that the multi-axis masking task successfully forces the architecture to learn robust, multi-modal features.

**Fig. 2:**
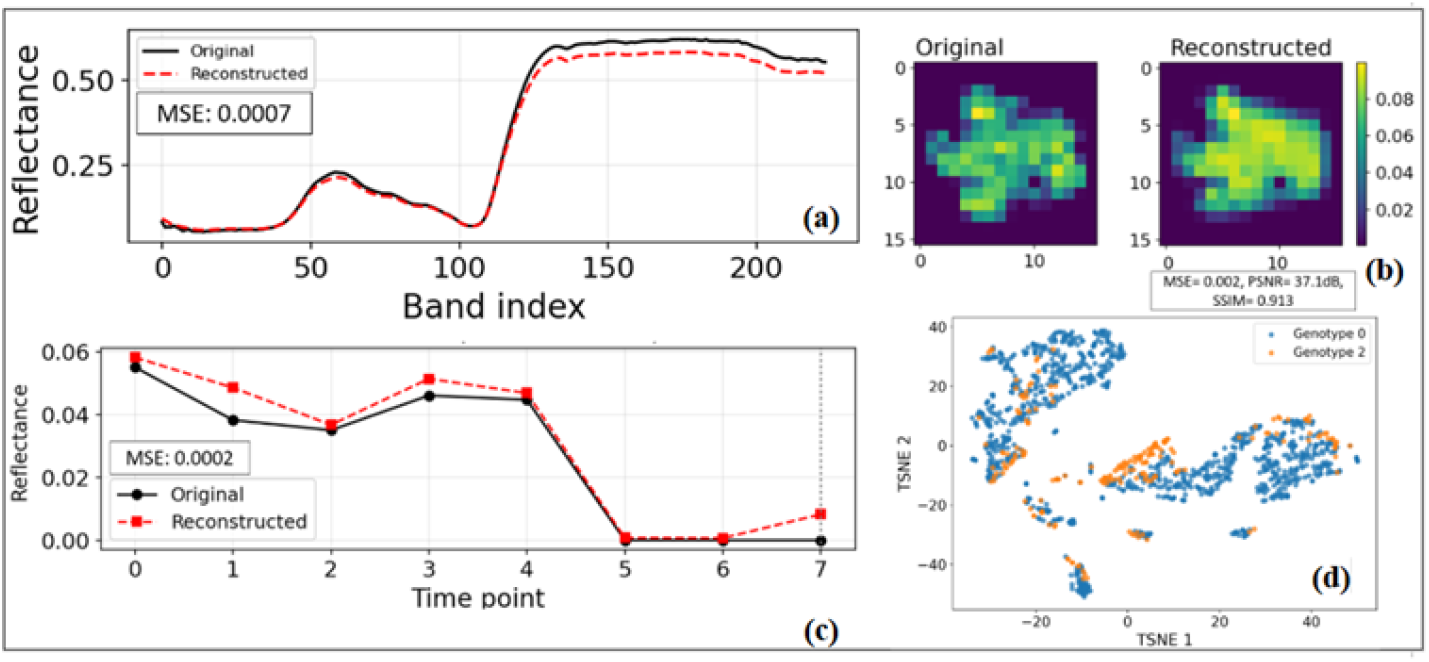
Pre-training reconstruction fidelity and global latent space topology. (a) Spectral reconstruction of a masked foreground patch. (b) Spatial reconstruction comparison across the 16 × 16 token grid. (c) Temporal trajectory reconstruction tracking plant reflectance over development. (d) 2D t-SNE projection of the global latent space for a representative target SNP.

To assess unsupervised feature-mapping, we visualized the global latent space using a 2D t-SNE projection of the mean-pooled temporal embeddings (**Fig. 2d**). The topology reveals a highly continuous, dispersed 2D landscape rather than a fragmented layout. This confirms that the pre-training task effectively coordinates multi-axis masking disruptions across the 3-layer transformer encoder. When mapping genetic variants for ant.loc9 across this latent space, genotype 2 (alternate allele, orange) organizes into distinct localized sub-clusters and directional branches embedded within the primary reference distribution (genotype 0, blue). This structural organization indicates that while global phenotypic traits govern the macro-topology, the encoder successfully preserves fine-grained, genotype-specific spectral-temporal variations. Downstream linear models can easily exploit these high-dimensional variations for accurate genotype prediction.

### 4.2 Model Performance on Functional SNPs

Our goal is representation learning rather than maximizing supervised performance. We therefore compare alternative feature representations using an identical downstream linear classifier (Logistic Regression), following standard self-supervised evaluation protocols [8, 22]. Performance was compared against two baseline frameworks: a raw spectral mean paired with Logistic Regression (Raw+LR) and PCA-reduced features paired with Logistic Regression (PCA+LR). SST-MAE achieved the highest classification performance across all four functional target loci, yielding the following AUC values: ant.loc5 = 0.91, ant.loc9 = 0.90, pale.loc4 = 0.88, and serr.loc5 = 0.77 (Fig. 3a; a representative run). Table 1 reports the mean *±* std across multiple runs. In contrast, the baseline configurations were comparatively lower, with performance metrics failing to exceed an AUC threshold of 0.85 across individual markers in both the PCA+LR pipeline (Fig. 3b) and the Raw+LR pipeline (Fig. 3c). The performance gap was most prominent for the serr.loc5 marker, where the SST-MAE architecture (Fig. 3a) outperformed the Raw+LR baseline model (Fig. 3c) by an absolute margin of +0.09 AUC points (0.77 vs. 0.68). This difference highlights the advantage of learned spatio-spectral-temporal features for structural plant characteristics that express as subtle morphological variants rather than macro-pigmentation differences readily captured by bulk averages. The baseline performance beneath the 0.85 AUC threshold demonstrates that SNP classification accuracy is affected by feature quality rather than backend classifier limitations. Self-supervised pretraining via SST-MAE successfully preserves genotype-specific structural patterns that linear projections fail to encode. The strong performance for anthocyanin loci likely reflects the direct influence of flavonoid biosynthesis on leaf reflectance within the visible spectrum [23]. In contrast, the lower performance for serr.loc5 is expected: leaf serration is a morphological trait that depends on developmental timing and requires longer temporal context before becoming distinguishable [24]. This confirms that SST-MAE captures biologically meaningful processes rather than memorizing spectral signatures.

**Table 1.**
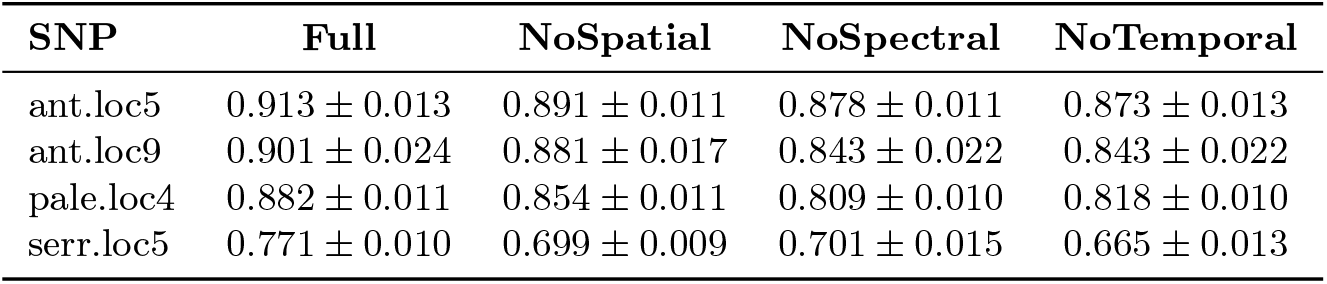
Ablation study results: Predictive performance (mean AUC std) of SST-MAE variants with spatial, spectral, or temporal masking omitted across four SNPs.

| SNP | Full | NoSpatial | NoSpectral | NoTemporal |
| --- | --- | --- | --- | --- |
| ant.loc5 | 0.913 ± 0.013 | 0.891 ± 0.011 | 0.878 ± 0.011 | 0.873 ± 0.013 |
| ant.loc9 | 0.901 ± 0.024 | 0.881 ± 0.017 | 0.843 ± 0.022 | 0.843 ± 0.022 |
| pale.loc4 | 0.882 ± 0.011 | 0.854 ± 0.011 | 0.809 ± 0.010 | 0.818 ± 0.010 |
| serr.loc5 | 0.771 ± 0.010 | 0.699 ± 0.009 | 0.701 ± 0.015 | 0.665 ± 0.013 |

**Fig. 3:**
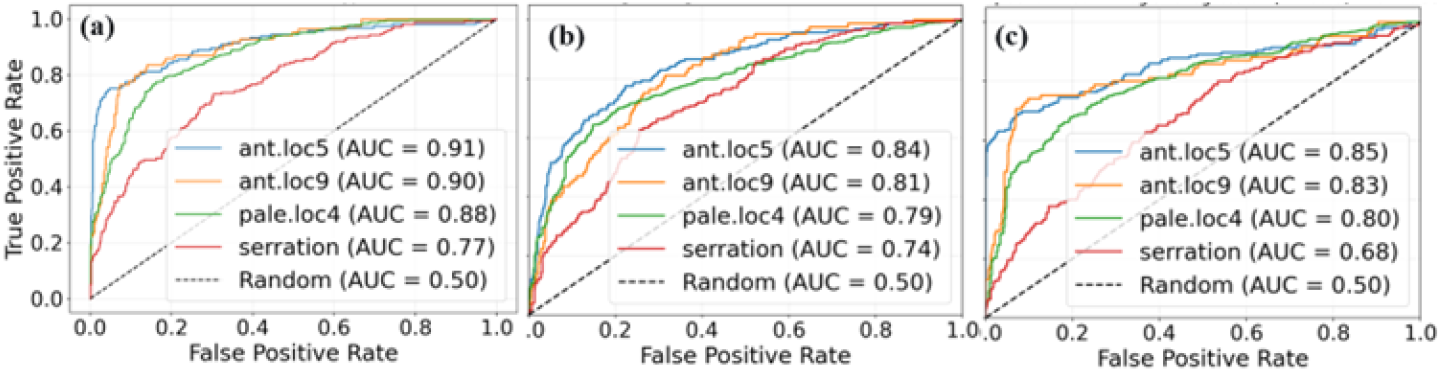
Performance comparison evaluated across three processing pipelines: (a) SST-MAE, (b) PCA+LR, and (c) raw+LR models.

### 4.3 Few-Shot Genotype Prediction

A key advantage of self-supervised pre-training such as SST-MAE is label efficiency, surpassing the need for annotated phenotypic labels. To evaluate the label efficiency of SST-MAE, we trained logistic regression classifiers on the learned latent representations using increasing proportions of labeled plants (10%, 30%, 50%, 70%, and 100%; see Supplementary Table S2 for absolute sample counts per SNP and partition). Label fractions below 10% were not evaluated; at 1–5% fractions the minority class (Class 2) would yield fewer than 40–80 labeled samples per SNP (based on Supplementary Tables S1 and S2), making reliable classifier training statistically unfeasible. The SST-MAE encoder was always pre-trained on the complete unlabeled dataset; only the amount of labeled data available for downstream classifier training was varied. This ensures that any improvements in label efficiency arise from the quality of the learned representations rather than from the amount of pre-training data. This was compared to the two baselines: Raw+LR and PCA+LR.

SST-MAE consistently outperformed both baselines at nearly most label fraction across the four functional SNPs. At the extreme low-label regime of 10% labels, SST-MAE achieved AUROCs of approximately 0.80 (Fig. 4a), 0.76 (Fig. 4b), 0.81 (Fig. 4c), and 0.63 (Fig. 4d) for ant.loc5, ant.loc9, pale.loc4, and serr.loc5, respectively. Although PCA+LR remained competitive for the anthocyanin-associated loci at low label fractions, SST-MAE consistently matched or exceeded its performance while maintaining a better performance over Raw+LR.

**Fig. 4:**
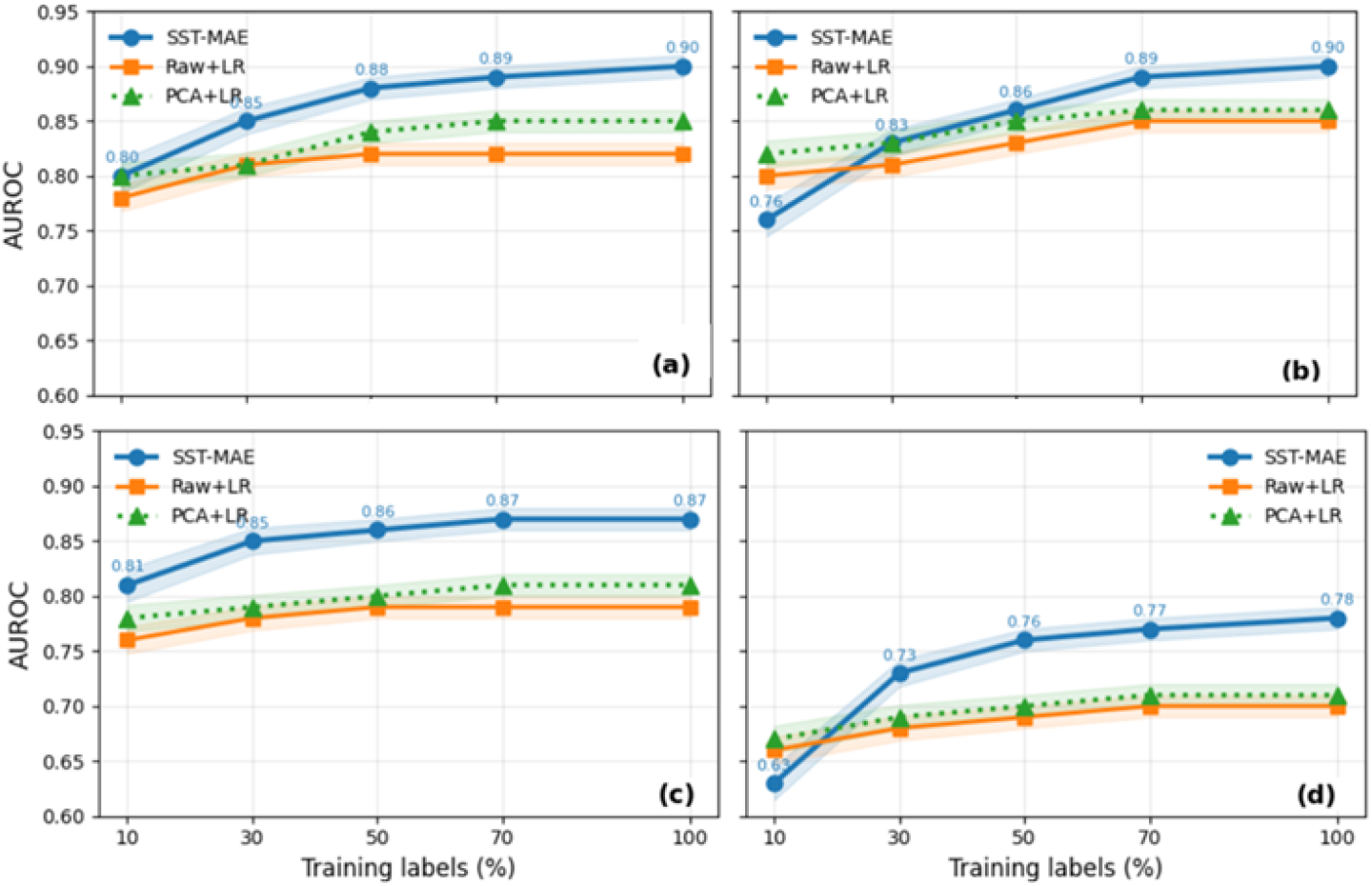
Few-shot learning curves (10%, 30%, 50%, 70%, 100% of labeled training plants; absolute counts in Supplementary Table S2) for four functional SNPs — (a) ant.loc5, (b) ant.loc9, (c) pale.loc4, and (d) serr.loc5 — comparing SST-MAE, Raw+LR, and PCA+LR models.

As the proportion of labeled data increased, the performance gap between SST-MAE and the baselines became more pronounced. For the anthocyanin loci (ant.loc5 and ant.loc9), PCA+LR approached SST-MAE at higher label fractions, suggesting that these pigmentation-related traits exhibit relatively strong linear spectral signatures that are partially captured by PCA. In contrast, SST-MAE demonstrated a substantially larger advantage for the serr.loc5 locus, where performance continued to improve with additional labeled data while both PCA+LR and Raw+LR plateaued. This observation indicates that the joint spatio-spectro-temporal representations learned by SST-MAE better capture the complex morphological characteristics associated with leaf serration than conventional linear feature representations. Together, these results demonstrate that the learned features are highly sample-efficient and predictive of genetic variation, proving valuable for capturing complex morphological variants where traditional linear baselines fall short.

### 4.4 Temporal Generalization

#### Cross-Time Transfer

To assess temporal generalization, we trained genotype classifiers on feature subsets extracted exclusively from the first three time points (*T*_1_–*T*_3_; early vegetative growth, 57–66 days after sowing) and evaluated them on the final two time points (*T*_7_–*T*_8_; late maturity, 99–106 days after sowing) (Fig. 5a). The framework maintained high AUROC on the early time points, with performance declining substantially for some traits when transferred to unseen late-stage data. The degree of transferability varied by trait. pale.loc4 exhibited the strongest temporal stability, decreasing only from 0.855 to 0.781 (*Δ*AUROC = 0.074). ant.loc5 showed a moderate decline from 0.906 to 0.751 (*Δ*AUROC = 0.155), while ant.loc9 dropped from 0.883 to 0.680 (*Δ*AUROC = 0.203). In contrast, serr.loc5 collapsed from 0.748 to 0.486 (*Δ*AUROC = 0.262), falling near chance on the late time points. This suggests that morphological traits like leaf serration are not only harder to predict overall, but also exhibit substantially poorer temporal generalization than biochemical traits, potentially because their developmental trajectories diverge more across genotypes during early growth.

**Fig. 5:**
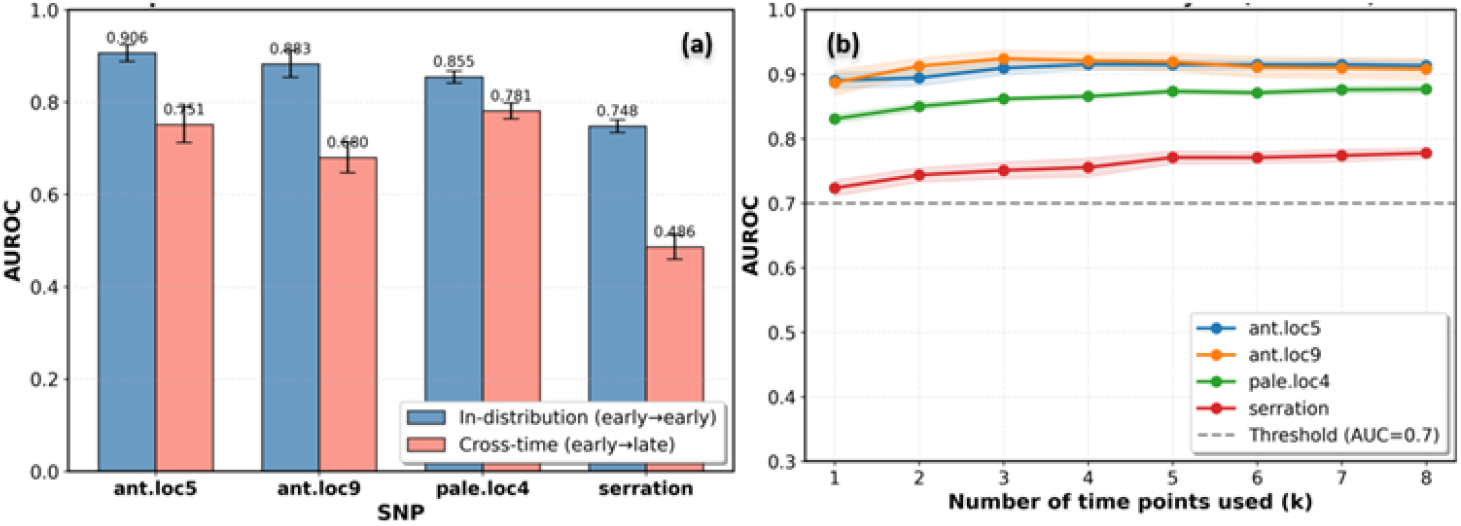
Temporal generalization analysis. (a) Cross-time transfer analysis showing how model trained on earlier time points can predict using later time points. (b) Time-to-detection time point showing when traits are detectable on the HSI data.

### Time-to-detection Analysis

To determine the earliest detectable genotype signal, performance was evaluated using progressively longer temporal sequences (*k* = 1–8; Fig. 5b). All four SNPs exceeded the detection threshold (AUROC ≥ 0.70) using only the first time point, demonstrating that SST-MAE encodes genotype-discriminative information at early vegetative stages. Performance improved as additional time points were added, with the majority of the gain achieved by *k* = 3–4 time points. The anthocyanin SNPs ant.loc5 and ant.loc9 achieved AUROC ≈0.89 at *k* = 1 and stabilized around 0.91 after three time points. pale.loc4 increased from 0.83 to 0.87, plateauing by *k* = 5, whereas serr.loc5 improved gradually from 0.72 to 0.78 by *k* = 8, indicating that structural traits require a longer temporal context than biochemical traits. All SNPs exceeded chance with the full time series, confirming that SST-MAE captures temporal dynamics. These results show that the detectability of genetic effects using HSI varies by trait: pigmentation traits may appear earlier and more prominent, while morphological traits require more temporal information or perhaps they may be difficult to capture intrinsically.

#### Ablation Study: Masking Modalities Contributions

To evaluate the individual contributions of each masking dimension, we systematically omitted spatial, spectral, and temporal masking blocks during pre-training and evaluated downstream classification performance (Table 1). The experimental results demonstrate that the full SST-MAE framework provides the most robust regularization, achieving the highest classification accuracy across nearly all target loci. For the anthocyanin-associated markers ant.loc5 and ant.loc9, the full model reached peak performance with AUCs of 0.913 and 0.901, respectively (Table 1). Removing the spatial masking module (NoSpatial) led to a drop in prediction accuracy across both pigmentation traits, proving that localized patch provides essential spatial regularization.

For pale.loc4, the performance remained stable between the full configuration (AUC = 0.882) and the NoSpatial variant (AUC = 0.854), with the difference falling well within the standard deviation range, while declining more noticeably when spectral masking was disabled (AUC = 0.809). This suggests that spectral signatures carry significant predictive weight for identifying internal leaf chlorosis patterns. The importance of the joint masking architecture is mostly reflected in the serr.loc5 trait. While individual ablation variants struggled— with the removal of temporal masking causing the sharpest decline to an AUC of 0.665—the Full SST-MAE model achieved a substantial performance, reaching an AUC of 0.771. Overall, the full SST-MAE exceeds random baselines (AUC ≥ 0.65), demonstrating its ability to learn biologically meaningful and transferable representations.

## 5 Conclusion

We presented SST-MAE, the first self-supervised framework to jointly model spatial, spectral, and temporal dimensions of plant hyperspectral time series, learning genotype-discriminative representations without phenotypic labels. By reconstructing masked information across all three axes, the framework captures developmental trajectories rather than static snapshots, and the frozen encoder transfers to downstream SNP classification with substantially better label efficiency than raw spectral or PCA-based baselines. Beyond raw performance, our analyses point to a more nuanced picture of what these representations capture: masking contributions are trait-dependent rather than uniformly beneficial, and temporal context matters most for morphological traits like leaf serration, which remain harder to predict than pigmentation loci and require longer developmental windows before becoming distinguishable. This suggests that genotype-dependent signals emerge at different rates and through different mechanisms during development — a distinction a static, single-time-point model cannot recover. SST-MAE offers plant breeders a scalable route to genotype screening from unlabeled hyperspectral collections, reducing reliance on manual phenotyping, and the framework should extend naturally to other crops and longitudinal imaging settings where labeled data remain scarce.

## Acknowledgements

This work is part of the LettuceKnow project (project number 1.2 of the research Perspective Program P19-17). The project is also partly sponsored by the Dutch Research Council (NWO), the associated Breeding companies: BASF, Bejo Zaden B.V., Limagrain, Enza Zaden Research & Development B.V., Rijk Zwaan Breeding B.V., Syngenta Seeds B.V., and Takii and Company Ltd., and the Foundation for Food and Agriculture Research (FFAR). The authors declare no competing interests.

## Notes

### Competing Interest Statement

The authors have declared no competing interest.

## References

1. A. Rasheed and X. Xia, “From markers to genome-based breeding in wheat,” Theor. Appl. Genet., vol. 132, no. 3, pp. 767–784, 2019.

2. R. Dijkhuizen, A. L. Van Eijnatten, S. L. Mehrem, and E. Van Den Bergh, “From aerial drone to quantitative trait locus: leveraging next-generation phenotyping to reveal the genetics of color and height in field-grown Lactuca sativa,” 2025, doi: 10.1111/tpj.70405.

3. T. Adão, J. Hruška, L. Pádua, J. Bessa, E. Peres, R. Morais, and J. J. Sousa, “Hyperspectral Imaging: A Review on UAV-Based Sensors, Data Processing and Applications for Agriculture and Forestry,” Remote Sens., vol. 9, no. 11, p. 1110, 2017, doi: 10.3390/rs9111110.

4. F. G. Okyere, D. Cudjoe, P. Sadeghi-Tehran, N. Virlet, A. B. Riche, M. Castle, L. Greche, F. Mohareb, D. Simms, M. Mhada, and M. J. Hawkesford, “Modeling the spatial-spectral characteristics of plants for nutrient status identification using hyperspectral data and deep learning methods,” Front. Plant Sci., vol. 14, p. 1209500, 2023, doi: 10.3389/fpls.2023.1209500.

5. S. Mehrem, A. Zijl, M. De Haan, G. Van Den Ackerveken, and B. L. Snoek, “Spectral Phenotyping Reveals Time-Specific QTLs in Field-Grown Lettuce,” 2026.

6. A. Kamilaris and F. X. Prenafeta-Boldú, “Deep learning in agriculture: A survey,” Comput. Electron. Agric., vol. 147, pp. 70–90, 2018.

7. Y. Cong, S. Khanna, C. Meng, P. Liu, E. Rozi, Y. He, M. Burke, D. B. Lobell, and S. Ermon, “SatMAE: Pre-training Transformers for Temporal and Multi-Spectral Satellite Imagery,” in Proc. NeurIPS, 2022.

8. K. He, X. Chen, S. Xie, Y. Li, P. Dollár, and R. Girshick, “Masked Autoencoders Are Scalable Vision Learners,” in Proc. IEEE/CVF Conf. Comput. Vis. Pattern Recognit. (CVPR), pp. 15979–15988, 2022, doi: 10.1109/CVPR52688.2022.01553.

9. Z. Tong, Y. Song, J. Wang, and L. Wang, “VideoMAE: Masked Autoencoders are Data-Efficient Learners for Self-Supervised Video Pre-Training,” in Proc. NeurIPS, 2022.

10. W. Zheng, Q. Zhang, J. He, B. Han, Q. Zhang, and D. Sun, “Comparative evaluation of SNP-weighted, Bayesian, and machine learning models for genomic prediction in Holstein cattle,” 2025.

11. H. Wang et al., “Cropformer: An interpretable deep learning framework for crop genomic prediction,” Plant Commun., vol. 6, no. 3, p. 101223, 2025.

12. A. Stylianou, R. Pless, N. Shakoor, and T. C. Mockler, “Classification and Visualization of Genotype × Phenotype Interactions in Biomass Sorghum,” in Proc. IEEE/CVF Int. Conf. Comput. Vis. Workshops (ICCVW), pp. 1352–1361, 2021.

13. P. Ballesta, C. Maldonado, F. Mora-Poblete, D. Mieres-Castro, A. del Pozo, and G. A. Lobos, “Spectral-Based Classification of Genetically Differentiated Groups in Spring Wheat Grown under Contrasting Environments,” Plants, vol. 12, no. 3, p. 440, 2023.

14. T. Gill, S. K. Gill, D. K. Saini, Y. Chopra, J. P. de Koff, and K. S. Sandhu, “A Comprehensive Review of High Throughput Phenotyping and Machine Learning for Plant Stress Phenotyping,” Phenomics, vol. 2, no. 3, pp. 156–183, 2022.

15. K. He, H. Fan, Y. Wu, S. Xie, and R. Girshick, “Momentum Contrast for Unsupervised Visual Representation Learning,” in Proc. IEEE/CVF Conf. Comput. Vis. Pattern Recognit. (CVPR), pp. 9729–9738, 2020.

16. H. Bao, L. Dong, S. Piao, and F. Wei, “BEiT: BERT Pre-Training of Image Transformers,” in Proc. Int. Conf. Learn. Represent. (ICLR), 2022.

17. L. Zhu, J. Wu, B. Wang, Y. Liao, and D. Gu, “SpectralMAE: Spectral Masked Autoencoder for Hyperspectral Remote Sensing Image Reconstruction,” Sensors, vol. 23, no. 7, p. 3728, 2023, doi: 10.3390/s23073728.

18. G. Bertasius, H. Wang, and L. Torresani, “Is Space-Time Attention All You Need for Video Understanding?,” in Proc. Int. Conf. Mach. Learn. (ICML), 2021.

19. T. Wei, R. van Treuren, X. Liu, Z. Zhang, J. Chen, Y. Liu, S. Dong, P. Sun, T. Yang, T. Lan, X. Wang, Z. Xiong, Y. Liu, J. Wei, H. Lu, S. Han, J. C. Chen, X. Ni, J. Wang, H. Yang, X. Xu, Y. Kuang, W. Wang, H. Zhang, J. D. G. Jones, K. Wang, and Z. Liu, “Whole-genome resequencing of 445 Lactuca accessions reveals the domestication history of cultivated lettuce,” Nat. Genet., vol. 53, no. 5, pp. 752–760, 2021.

20. D. J. M. van Workum, S. L. Mehrem, B. L. Snoek, M. C. Alderkamp, D. Lapin, F. F. M. Mulder, G. Van den Ackerveken, and D. de Ridder, “Lactuca super-pangenome reduces bias towards reference genes in lettuce research,” BMC Plant Biol., vol. 24, no. 1, 2024.

21. A. Dosovitskiy, L. Beyer, A. Kolesnikov, D. Weissenborn, X. Zhai, T. Unterthiner, M. Dehghani, M. Minderer, G. Heigold, S. Gelly, J. Uszkoreit, and N. Houlsby, “An Image Is Worth 16x16 Words: Transformers for Image Recognition at Scale,” in Proc. Int. Conf. Learn. Represent. (ICLR), 2021.

22. T. Chen, S. Kornblith, M. Norouzi, and G. Hinton, “A Simple Framework for Contrastive Learning of Visual Representations,” in Proc. Int. Conf. Mach. Learn. (ICML), pp. 1597–1607, 2020.

23. A. A. Gitelson, M. N. Merzlyak, and O. B. Chivkunova, “Optical properties and nondestructive estimation of anthocyanin content in plant leaves,” Photochem. Photobiol., vol. 74, no. 1, pp. 38–45, 2001.

24. I. Rubio-Somoza, C. M. Zhou, A. Confraria, C. Martinho, P. von Born, A. Baena-Gonzalez, J. Wang, and D. Weigel, “Temporal control of leaf complexity by miRNA-regulated licensing of protein complexes,” Curr. Biol., vol. 24, no. 22, pp. 2714–2719, 2014.

