## supplementary information for "SST-MAE: Learning Spectral-Spatio-Temporal Representations from Plant Hyperspectral Time Series to Discover Complex Genotype-Phenotype Relations"

### SST-MAE: Supplementary Material


#### Supplementary Material

**Table 1: Table S1.** Sample sizes and class distributions for the four SNPs. The 80/20 genotype-level split yields 11,376 training and 2,880 test plant images. Heterozygotes are included in total counts but excluded from binary class columns.

| SNP | Total samples | Training (n) | Test (n) | Class 0 (train) | Class 2 (train) | Class 0 (test) | Class [illegible] |
|---|---|---|---|---|---|---|---|
| ant.loc5 | 14,256 | 11,376 | 2,880 | 10,152 | 1,224 | 1,656 | [illegible] |
| ant.loc9 | 14,256 | 11,376 | 2,880 | 10,512 | 792 | 2,376 | [illegible] |
| pale.loc4 | 14,256 | 11,376 | 2,880 | 4,824 | 6,192 | 1,368 | [illegible] |
| serr.loc5 | 14,256 | 11,376 | 2,880 | 9,360 | 2,016 | 2,088 | [illegible] |

**Table 2: Table S2.** Few-shot training set sizes (10–100%) per SNP. Fractions are based on the total homozygous training pool. The fixed test set (held-out genotypes) remains fully labeled.

| SNP | 10% | 30% | 50% | 70% | 100% | Fixed test set |
|---|---|---|---|---|---|---|
| ant.loc5 | 1,138 | 3,413 | 5,688 | 7,963 | 11,376 | 2,880 |
| ant.loc9 | 1,130 | 3,391 | 5,652 | 7,913 | 11,304 | 2,808 |
| pale.loc4 | 1,102 | 3,305 | 5,508 | 7,711 | 11,016 | 2,880 |
| serr.loc5 | 1,138 | 3,413 | 5,688 | 7,963 | 11,376 | 2,880 |

**Table 3: Table S3.** Computational cost summary for SST-MAE pre-training and inference.

| **Metric** | **Value** |
|---|---:|
| GPU | NVIDIA RTX A6000 (48 GB) |
| Training epochs | 100 (early stopping at 72) |
| Batch size | 16 |
| Total parameters | 4.2 million |
| Encoder parameters | 2.8 million |
| Decoder parameters | 1.4 million |
| Pre-training time | $\approx$8.5 hours |
| Inference (feature extraction) per plant | $\approx$45 ms |
